# Characterization of young and old adult brains: An EEG functional connectivity analysis

**DOI:** 10.1101/495564

**Authors:** Bahar Moezzi, Latha Madhuri Pratti, Brenton Hordacre, Lynton Graetz, Carolyn Berryman, Louise M. Lavrencic, Michael C. Ridding, Hannah A. D. Keage, Mark D. McDonnell, Mitchell R. Goldsworthy

**Author notes:** Corresponding author (Bahar Moezzi). Equal contributors (Bahar Moezzi, Latha Madhuri Pratti).

## Abstract

Brain connectivity studies have reported that functional networks change with older age. We aim to (1) investigate whether electroencephalography (EEG) data can be used to distinguish between individual functional networks of young and old adults; and (2) identify the functional connections that contribute to this classification. Two eyes-open resting-state EEG recording sessions with 64 electrodes for each of 22 younger adults (19-37 years) and 22 older adults (63-85 years) were conducted. For each session, imaginary coherence matrices in theta, alpha, beta and gamma bands were computed. A range of machine learning classification methods were utilized to distinguish younger and older adult brains. A support vector machine (SVM) classifier was 94% accurate in classifying the brains by age group. We report decreased functional connectivity with older age in theta, alpha and gamma bands, and increased connectivity with older age in beta band. Most connections involving frontal, temporal, and parietal electrodes, and approximately two-thirds of connections involving occipital electrodes, showed decreased connectivity with older age. Just over half of the connections involving central electrodes showed increased connectivity with older age. Functional connections showing decreased strength with older age had significantly longer electrode-to-electrode distance than those that increased with older age. Most of the connections used by the classifier to distinguish participants by age group belonged to the alpha band. Findings suggest a decrease in connectivity in key networks and frequency bands associated with attention and awareness, and an increase in connectivity of the sensorimotor functional networks with ageing during a resting state.

## 1. Introduction

Advanced age is associated with a progressive decline in cognition, particularly in the domains of attention, memory and executive function (Hedden & Gabrieli, 2004; Raz & Rodrigue, 2006). While there has been considerable research investigating the pathophysiological changes that characterize Alzheimer’s disease and other age-related neurodegenerative disorders (Goedert & Spillantini, 2006; Y. Huang & Mucke, 2012), much less is known about the neural processes affecting cognition in normal, non-pathological ageing. A greater understanding of the basic neurophysiology of healthy human ageing may prove critical for devising novel therapeutic approaches for slowing or reversing age-related cognitive decline.

The long-range connectivity of resting-state brain networks decreases in healthy ageing (Esposito et al., 2008; Ferreira & Busatto, 2013; Hafkemeijer, van der Grond, & Rombouts, 2012; Koch et al., 2010; Meunier, Achard, Morcom, & Bullmore, 2009; Mevel, Chételat, Eustache, & Desgranges, 2011; Tomasi & Volkow, 2012). Using fMRI age-related decreases in resting-state functional connectivity have been observed in the default mode network (DMN) and the dorsal attention network (DAN) (Hafkemeijer et al., 2012; Mevel et al., 2011), both of which are heavily implicated in attention, memory and executive functions (van Den Heuvel & Hulshoff Pol, 2010). The DMN comprises a set of regions active at rest, including the medial prefrontal cortex, the inferior parietal lobule, the hippocampus and the posterior cingulate cortex/retrosplenial cortex/precuneus (Buckner, Andrews-Hanna, & Schacter, 2008). The DAN includes a set of brain regions comprising the prefrontal, anterior cingulate and posterior parietal cortices (Tomasi & Volkow, 2012).

It is suggested that age-related changes in the motor networks may be different from those in the DMN and the DAN. Using fMRI increased functional connectivity has been found in motor and subcortical networks in healthy ageing (Tomasi & Volkow, 2012). In a resting-state functional magnetic resonance imaging (fMRI) study of 913 healthy adults from the 1000 Functional Connectomes Project repository, Tomasi and Volkow (2012) observed that advancing age was associated with increased functional connectivity in the somatosensory and motor cortices, cerebellum and brainstem. However, other studies have reported reverse findings (Allen et al., 2011; Toussaint et al., 2011; T. Wu et al., 2007).

EEG is a non-invasive method of recording cortical activity through the scalp (Niedermeyer & da Silva, 2005). While the spatial resolution of EEG is limited compared to fMRI, its high temporal precision makes it possible to measure neural oscillations that occur in the millisecond time range due to synchronized rhythmical firing of populations of neurons (Niedermeyer & da Silva, 2005). This oscillatory activity is thought to play a key role in coordinating activity in large-scale brain networks, facilitating information flow between distributed brain regions when oscillations are synchronized between regions (Siegel, Donner, & Engel, 2012). Neural oscillations are grouped into specific frequency bands, each reflecting neural activity within particular brain structures and under different brain states. For instance, theta band oscillations have been linked to hippocampal-dependent behaviours such as spatial navigation and learning (Araújo, Baffa, & Wakai, 2002; Kahana, Sekuler, Caplan, Kirschen, & Madsen, 1999), and are also increased in frontal and parietal brain regions during different types of working memory and attentional control tasks (Klimesch, Freunberger, Sauseng, & Gruber, 2008; Sauseng, Griesmayr, Freunberger, & Klimesch, 2010). Alpha band oscillations are dominant in occipital regions, particularly during relaxed wakefulness with eyes closed, and are also thought to play an important role in attention and working memory by gating sensory processing to protect information held online from sensory interference (Klimesch, Sauseng, & Hanslmayr, 2007). Beta band oscillations have been linked to sensorimotor network activity (Pfurtscheller, Stancak Jr, & Neuper, 1996; Roopun et al., 2006). Gamma band oscillations, on the other hand, have been associated with a range of sensory and cognitive processes (Başar, Başar-Eroglu, Karakaş, & Schürmann, 2001; Fries, 2009), although appear particularly affected by muscle and electrical noise (Pope, Fitzgibbon, Lewis, Whitham, & Willoughby, 2009; Whitham et al., 2007).

Functional connectivity can be estimated in specific frequency bands using EEG recordings (Sakkalis, 2011). Frequency specific changes in functional connectivity throughout the lifespan have been previously reported (Micheloyannis et al., 2009; Smit et al., 2012). Smit et al. (2012) investigated the change in the functional brain connectivity from ages 5 through 71 years using resting-state EEG. Smit et al. (2012) reported large increases in theta, alpha and beta functional connectivity from childhood to adolescence that continued at a slower pace into adulthood (peaking at 50 years) and decreases in theta, alpha and beta functional connectivity above 50 years. Micheloyannis et al. (2009) used resting-state and task (mathematical thinking) EEG from twenty children and twenty young adults and reported lower beta and gamma connectivity in both rest and during task in adults. Vysata et al. (2014) used resting-state EEG in a group of 17,722 healthy professional drivers with a mean age of 43.2 years (SD = 11.2 years) and reported average functional connectivity over the whole scalp increased with older age in the beta band and decreased with older age in theta and alpha bands. However, it is still unknown how frequency-specific functional connections change with older age

Machine learning classifiers (Pereira, Mitchell, & Botvinick, 2009) provide a powerful approach to investigate age-related differences in functional networks. One of the commonly used machine learning classifiers is support vector machine (SVM). A crucial aspect of SVM is that it identifies the features that drive classifier performance. An SVM algorithm trained on a training dataset can generate feature weights corresponding to the relative contribution of an individual feature to successful differentiation of the two groups. Following this, the classifier can be applied to a separate testing dataset to assess the accuracy of the classifier in differentiating the two groups. SVM classifiers have been used to classify individual brains by chronological age using resting-state fMRI connectivity data. Dosenbach et al. (2010) used data from 238 fMRI scans of participants aged 7 to 30 years old. They classified children and adult brains with 91% accuracy, which was replicated as 92% and 93% accuracy with two other datasets comprising of 195 and 186 scans. Meier et al. (2012) used three scans from 26 younger adults (18-35 years old) and 26 older adults (55-85 years old) and were able to classify younger and older adult brains with 84% accuracy. They showed that the majority of the functional connections that distinguished older and younger adults came from regions belonging to the sensorimotor and cingulo-opercular functional networks. They reported a decrease in long-range functional connectivity and an increase in short-range functional connectivity with older age. Machine learning has been successfully applied to resting-state EEG data in the classification of typically developing children from a group of infants at high risk for autism spectrum disorder (Kousarrizi, Ghanbari, Gharaviri, Teshnehlab, & Aliyari, 2009) and in the classification of alcoholics and non-alcoholics (Bosl, Tierney, Tager-Flusberg, & Nelson, 2011). Investigating EEG functional connectivity characteristics of healthy ageing will provide further understanding of the physiological processes that mediate changes in functional connectivity as humans age.

We aim at uncovering whether an individual resting EEG session can be used to distinguish young and old brains and most importantly, the identity of the functional connections that contribute to such classification. We hypothesized that long-range functional connectivity of theta and alpha band oscillations, particularly in frontal, parietal and occipital regions, would be decreased with older age.

## 2. Materials and Methods

### 2.1 Procedure

Resting-state EEG data from 22 younger adults aged 19 to 37 years (mean 24.3 years, SD 6.1 years, 9 male, 21 self-reported right-handed) and 22 older adults aged 63 to 85 years (mean 71 years, SD 6 years, 9 male, 22 right-handed according to the Flinders Handedness Survey (Nicholls, Thomas, Loetscher, and Grimshaw (2013)) were recorded. In line with Meier et al. (2012) we used more than one recording session from each participant. Our dataset consists of two separate resting-state recording sessions for each participant, recorded at least 7 days apart. The EEG recordings were baseline measurements in pre-intervention or sham conditions embedded in larger studies on the effects of transcranial magnetic stimulation and/or Noxious stimulation on brain dynamics. The EEG data for younger adults is published elsewhere in the context of a graph theoretical analysis (Moezzi, Hordacre, Berryman, Ridding, & Goldsworthy, 2018). Participants gave written informed consent in accordance with the World Medical Association Declaration of Helsinki to participate in this study. Ethical approval was provided by the University of Adelaide Human Research Ethics Committee and University of South Australia’s Human Research Ethics Committee.

### 2.2 EEG acquisition and pre-processing

We acquired three minutes of continuous resting-state EEG data (eyes open) using an ASA-lab EEG system (ANT Neuro, Enschede, Netherlands) or a TMSi EEG system (Twente Medical Systems International B.V, Oldenzaal, The Netherlands) using a Waveguard™ original cap with 64 sintered Ag-AgCl electrodes in standard 10-10 positions. During recording, participants were instructed to view a fixation point, remain still, quiet and relaxed to avoid blinking too much. Each participant’s two sessions were performed using the same EEG system. Signals were sampled at 2048 Hz, amplified 20 times, filtered (high pass, DC; low pass 553 Hz) and referenced to the average of all electrodes. Impedance was kept below 5 kΩ and the recorded data were stored on a computer for offline analysis.

EEG data were exported to MATLAB 9.0 (MathWorks, Inc., Natick, MA) for pre-processing and analysis. The EEG signals were segmented into epochs of 1 second. The baseline means were removed from the EEG dataset. Channels Fp1, Fpz, Fp2, M1, M2, AF7 and AF8 and channels that were disconnected during recording or dominated by exogenous artefact noise were removed. The data were filtered using a hamming windowed sinc finite impulse response filter (1-45 Hz). We excluded epochs contaminated by excessive deflection identified by a threshold for the maximum allowed amplitude for the EEG signals >100 µV (7% of epochs were excluded). Fast ICA artefact correction was implemented to correct for non-physiological artefacts (e.g. eye blinks and scalp muscle activity) (Delorme & Makeig, 2004). Missing channels were interpolated using super-fast spherical interpolation.

### 2.3 Power

We used the power spectra to set the range of the theta, alpha, beta and gamma frequency bands separately for younger and older adults. We computed power-spectra by performing time-frequency analysis on EEG time series over multiple trials using the multitaper method based on Hanning tapers. The analysis windows were centred in each trial at 0.01 s intervals with three cycles per time window. In line with (Haegens, Cousijn, Wallis, Harrison, & Nobre, 2014; Moretti et al., 2011), our alpha frequency was defined as the biggest local maximum within the extended range (5–14 Hz). The theta/alpha transition frequency (TF) was computed as the minimum power in the alpha frequency range. Theta band was defined from 4 Hz to TF. The gamma frequency was defined as the highest peak with frequency above 30 Hz (Herrmann, Lenz, Junge, Busch, & Maess, 2004) and up to 45 Hz (Hordacre, Moezzi, & Ridding, 2018). The beta band was defined as the frequencies between the higher endpoint of the alpha band and 30 Hz (Henelius, Korpela, & Huotilainen, 2011). We utilized FieldTrip which is a MATLAB software toolbox for EEG and MEG analysis to compute power (Oostenveld & Praamstra, 2001).

### 2.4 Functional connectivity

We constructed the functional connectivity matrices for each participant using imaginary coherence in theta, alpha, beta and gamma frequency bands. The *l*-th segment of the *i*-th time course is denoted by *x*_*i,l*_ and its Fourier transform by *X*_*i,l*_. The cross-spectral matrix is defined as

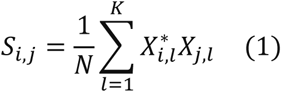

where (.)* denotes complex conjugation and *N* denotes the total number of segments. The coherency between the *i*-th and *j*-th times series is defined as

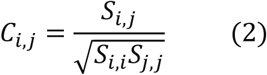

We computed the absolute value of imaginary coherence between each two EEG electrodes *m* and *n* as

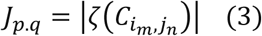

where ζ(.) denotes the imaginary part. Finally, matrix *J* was divided by its standard deviation using the Jackknife method to generate a connectivity matrix. To compute functional connectivity, we utilized the FieldTrip software package (Oostenveld & Praamstra, 2001).

### 2.5 Classification

Soft-margin SVM classification was performed using Scikit-Learn, a machine learning toolbox implemented in Python (Pedregosa et al., 2011). Local Outlier Factor and Isolation Forest algorithms were used to identify outliers (Breunig, Kriegel, Ng, & Sander, 2000; Liu, Ting, & Zhou, 2008). In line with previous literature, we mapped the data in higher-dimension using a radial basis function (RBF) as the underlying kernel and discriminated functional connectivity data as belonging to the younger or older adult group using a linear decision function (Meier et al., 2012). A leave-one-out cross-validation (LOOCV) procedure was performed to (1) identify the most significant features, (2) tune the hyper-parameters (C-regularization factor and gamma-kernel coefficient) of SVM and (3) determine the accuracy of the classifier. For each iteration of LOOCV, both sessions of a participant were removed, and the top 300 features (functional connections) were selected using two-sample t-tests on the training set and ranked according to their absolute t-statistics. We used the resulting t-scores as our feature weights. We kept 300 features because this was the approximate number of features that remained significant following False Discovery Rate correction. After selecting the top features, a nested LOOCV was performed to train the classifier and tune the hyperparameters. In the outer LOOCV loop, both the sessions of a participant were set aside for validating the classifier. The sessions of the remaining participants were used in the inner LOOCV loop to tune the hyperparameters of the classifier. Once the classifier is trained, the left-out sessions were separately classified as belonging to the younger or older adult group. The total accuracy of the classifier was determined by the percent of correctly classified sessions across all iterations. Receiver operating characteristics (ROC) curve was considered to calculate the area under curve score for a dataset having balanced classes (Fawcett, 2006). ROC takes into consideration the value of the threshold that identifies which class the session belongs to. A perfect classifier has a score of 1 whereas a classifier operating at random has a score of 0.5. For each group, young and old, the precision of the classifier was determined as the number of true positives divided by the sum of true positives and false positives. The recall of the classifier was determined by the number of true positives divided by the sum of true positives and false negatives.

Due to equal number of older and younger adults, the chance performance of the classifier generates an accuracy of 50%. In line with previous literature, we considered each iteration of the LOOCV as a Bernoulli trial with success probability of 0.5 (Pereira et al., 2009). By comparing the number of true positives to the number of sessions to be classified we also determined the probability of the accuracy occurring by chance.

In addition to SVM, we performed classification using alternative methods. Once the most significant features were identified, the LOOCV procedure was performed to tune the hyperparameters and determine the accuracy of each of the following methods: K-nearest neighbours (Cover & Hart, 1967) was performed by tuning the number of nearest neighbours, least squares linear classifier (Bishop, 2006), SVM classifier with linear kernel by tuning the regularization factor (C), and extreme learning machine (ELM) (Akusok, Bjork, Miche, & Lendasse, 2015; G.-B. Huang, Zhu, & Siew, 2004) with linear, RBF and sigmoid kernels by tuning the number of neurons to be added to the network.

### 2.6 Brain regions

We categorized electrodes into groups approximating frontal, central, parietal, temporal, and occipital brain regions based on an EEG study by Kamarajan et al. (2015) – see Figure 1. To compute the contribution of each region to the SVM classification, if a feature consisted of two electrodes from different brain regions, then half of the feature weight was assigned to each brain region. If both electrodes of a feature were from the same region, then the full feature weight was assigned to that region. The percentage contribution of each region was then normalized to account for differences in the total number of electrodes included in each region.

**Figure 1.**
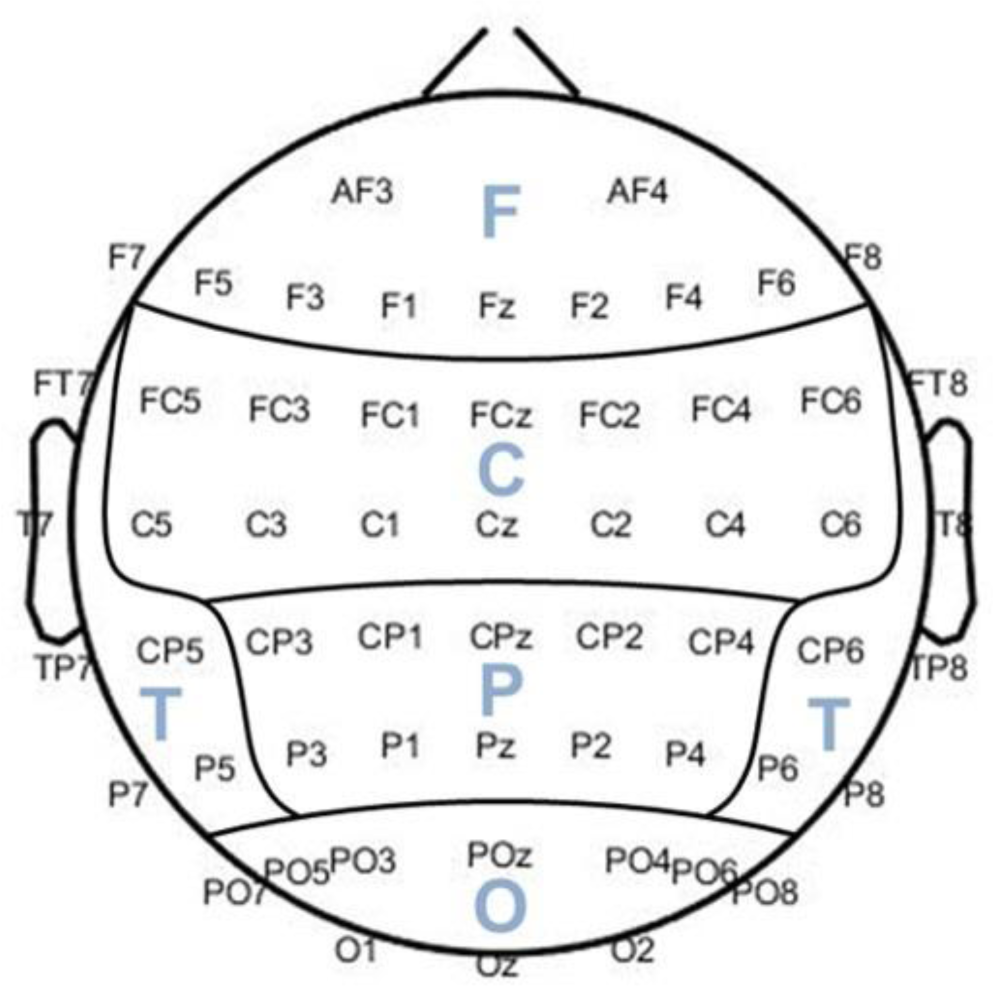
Grouping electrodes to approximate frontal (F), central (C), temporal (T), parietal (P), and occipital (O) regions.

### 2.7 Electrode-to-electrode distance

We characterized consensus features by the Euclidian distance between electrode pairs to investigate patterns of age-related differences. The 3-D electrode template we used was a standard 10-5 template constructed by Oostenveld and Praamstra (2001) electrode positions represented in millimetre in the Montreal Neurological Institute (MNI) coordinate system.

### 2.8 Statistical analysis

We compared the connections that decreased with older age to those that increased with older age based on electrode-to-electrode Euclidian distance using Mann-Whitney *U*-test. P-values < 0.05 were considered statistically significant.

## 3. Results

### 3.1 Power

We plotted the power spectra for younger and older adults for theta, alpha, beta and gamma bands (see Figure 2). For older adults, frequency ranges of 4-6 Hz and 7-13 Hz were selected for theta and alpha bands, respectively, whereas for younger adults, 4-7 Hz and 8-13 Hz were used. We set beta band to 14-30 Hz and gamma band to 31-45 Hz for both younger and older adults. While we used the power spectra to set the frequency bands we continue with analysing on functional connectivity data. Our main goal is to identify the distinguishing age-related differences in functional connectivity.

**Figure 2.**
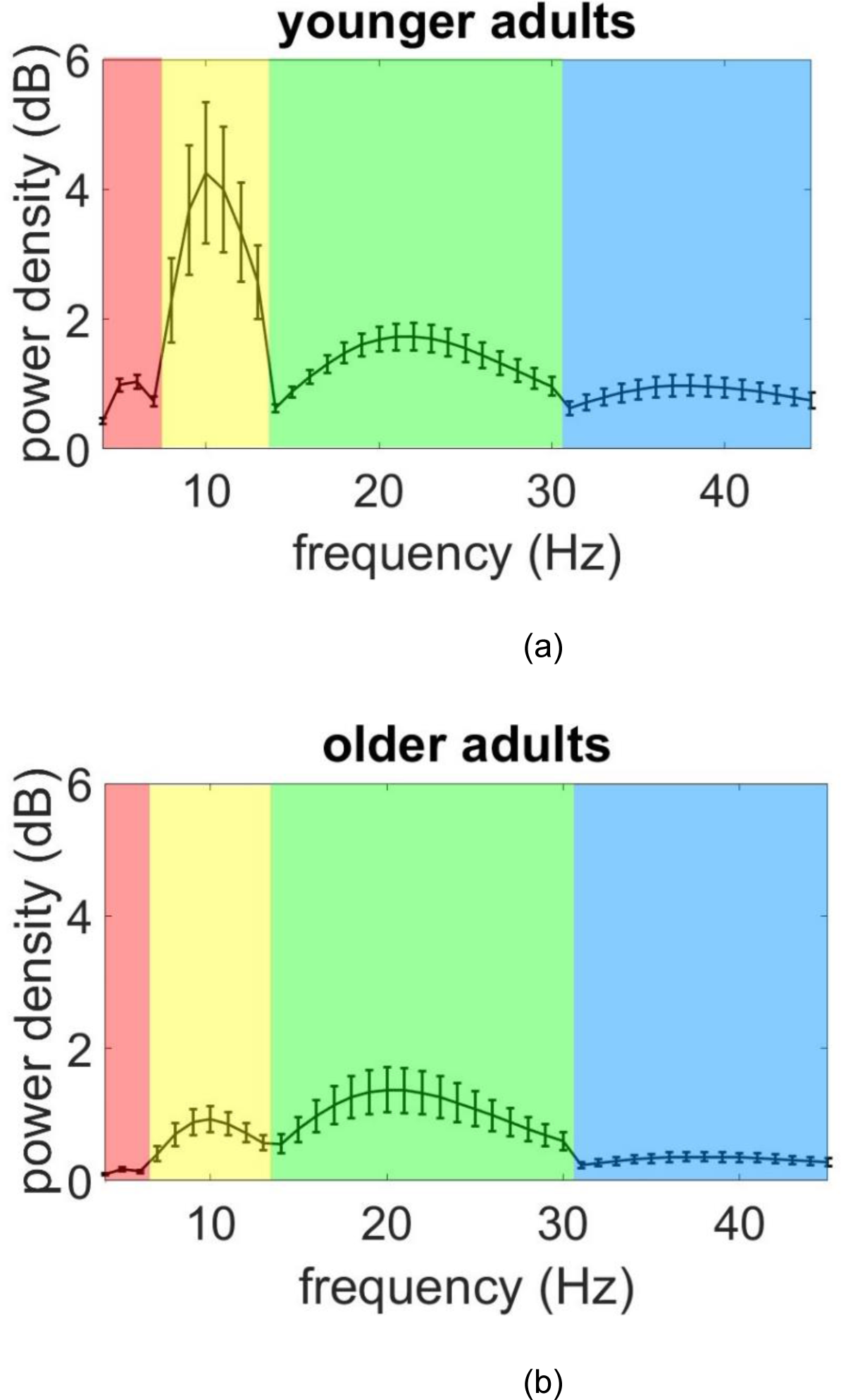
Mean Power spectra over all electrodes for (a) younger and (b) older adults. Error bars are SE. Red shade denotes the theta band, yellow shade denotes the alpha band, green shade denotes beta band and blue shade denote the gamma band.

### 3.2 Outliers

Of the 88 separate EEG recording sessions, Local Outlier Factor algorithm identified seven outliers, four of which belonged to the younger adult group and three to the older adult group. These same outliers were detected using Isolation Forest algorithm and therefore were removed from the dataset prior to SVM classification. To maintain equal numbers of sessions for both younger and older groups, we also randomly selected a session from the older adult group and removed it from the dataset.

### 3.3 Classifier performance

The classifier successfully classified younger adults from older adults with an accuracy of 94%. The 94% accuracy of the classifier was significantly higher than the accuracy by chance performance of the classifier (p < 10^−8^). Five recording sessions belonging to the older adult group were misclassified with one older adult having both sessions misclassified.

The accuracy achieved by the SVM (linear kernel), k-nearest neighbours, ELM (linear kernel), ELM (RBF kernel), ELM (sigmoid kernel) and least squares linear classifier was below 84%. The accuracy, area under ROC curve, precision and recall values for all the methods are summarized in Table 1. The SVM classifier with RBF kernel has the highest accuracy, area under ROC curve, precision and recall among all the methods tested.

**Table 1.** The accuracy of the different classification methods. SVM: support vector machine, ELM: extreme learning machine, RBF: radial basis function, ROC: receiver operating characteristics.

| Method | Accuracy | Area under ROC curve | Young |  | Old |  |
| --- | --- | --- | --- | --- | --- | --- |
|  |  |  | Precision | Recall | Precision | Recall |
| SVM (RBF kernel) | 94% | .97 | .89 | 1.0 | 1.0 | .88 |
| SVM (linear kernel) | 80% | .89 | .85 | .73 | .76 | .88 |
| K-nearest neighbours | 78% | .82 | .79 | .75 | .76 | .80 |
| ELM (linear kernel) | 70% | .80 | .70 | .70 | .70 | .70 |
| ELM (RBF kernel) | 80% | .78 | .82 | .78 | .79 | .83 |
| ELM (sigmoid kernel) | 84% | .84 | .87 | .80 | .81 | .88 |
| Least squares linear classifier | 68% | .75 | .68 | .68 | .69 | .68 |

### 3.4 Features

The features were analysed to uncover the patterns of functional connectivity changes that contributed to the SVM classification. From approximately 300 features obtained from each iteration of LOOCV, 162 features were identified to be common among all iterations (consensus features) – see Supplementary material Table 2. Eighty three percent of total consensus feature weights came from pairs of electrodes that showed lower functional connectivity in older adults than younger adults.

#### 3.4.1 Frequency bands

We further analysed the consensus features belonging to theta, alpha, beta and gamma frequency bands. The frequency band with the greatest amount of consensus feature weight associated with it was the alpha band (112 of 162 consensus features), followed by beta (28 features), theta (17 features), and gamma (5 features) (Figure 3). The consensus features belonging to theta, alpha and gamma bands had positive weights showing significantly higher functional connectivity in younger adults than in older adults. Consensus features of the beta band had negative weights showing significantly higher functional connectivity in older adults than younger adults.

**Figure 3.**
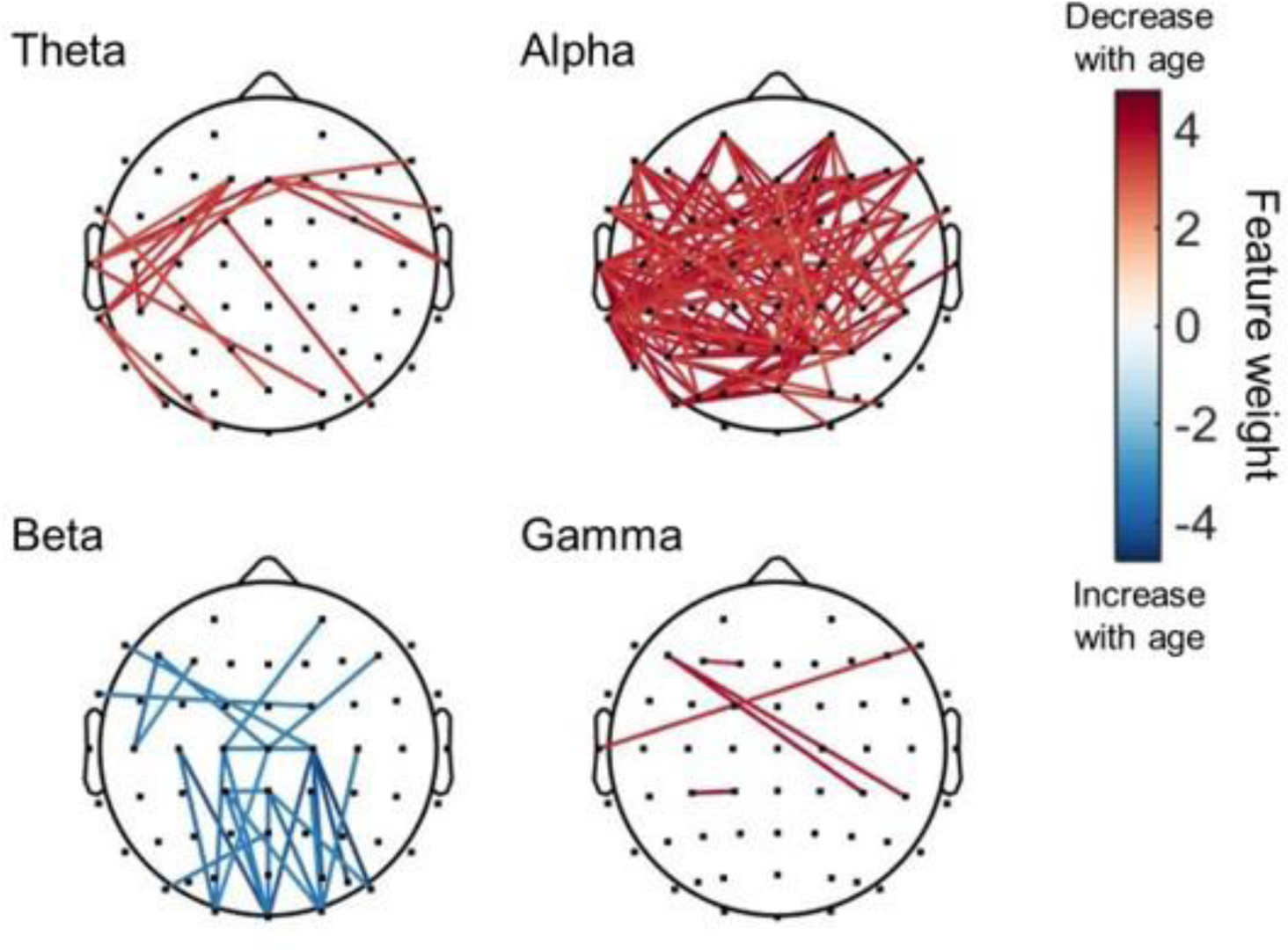
Illustration of the consensus features with decreased weight with older age (red) and the consensus features with increased weight with older age (blue) in theta, alpha, beta and gamma frequency bands.

#### 3.4.2 Brain regions

Electrodes were grouped as approximating five brain regions (temporal, frontal, occipital, parietal, central). The single electrode with the greatest amount of consensus feature weight associated with it was TP7 (temporal) followed by T7 (temporal), Pz (parietal), P2 (parietal) and POz (occipital). As seen in Figure 4, the large majority of features involving frontal, temporal, and parietal electrodes, and approximately two-thirds of features involving occipital electrodes, showed decreased connectivity with older age. Conversely, slightly more than half of the features involving central electrodes showed increased connectivity with older age.

**Figure 4.**
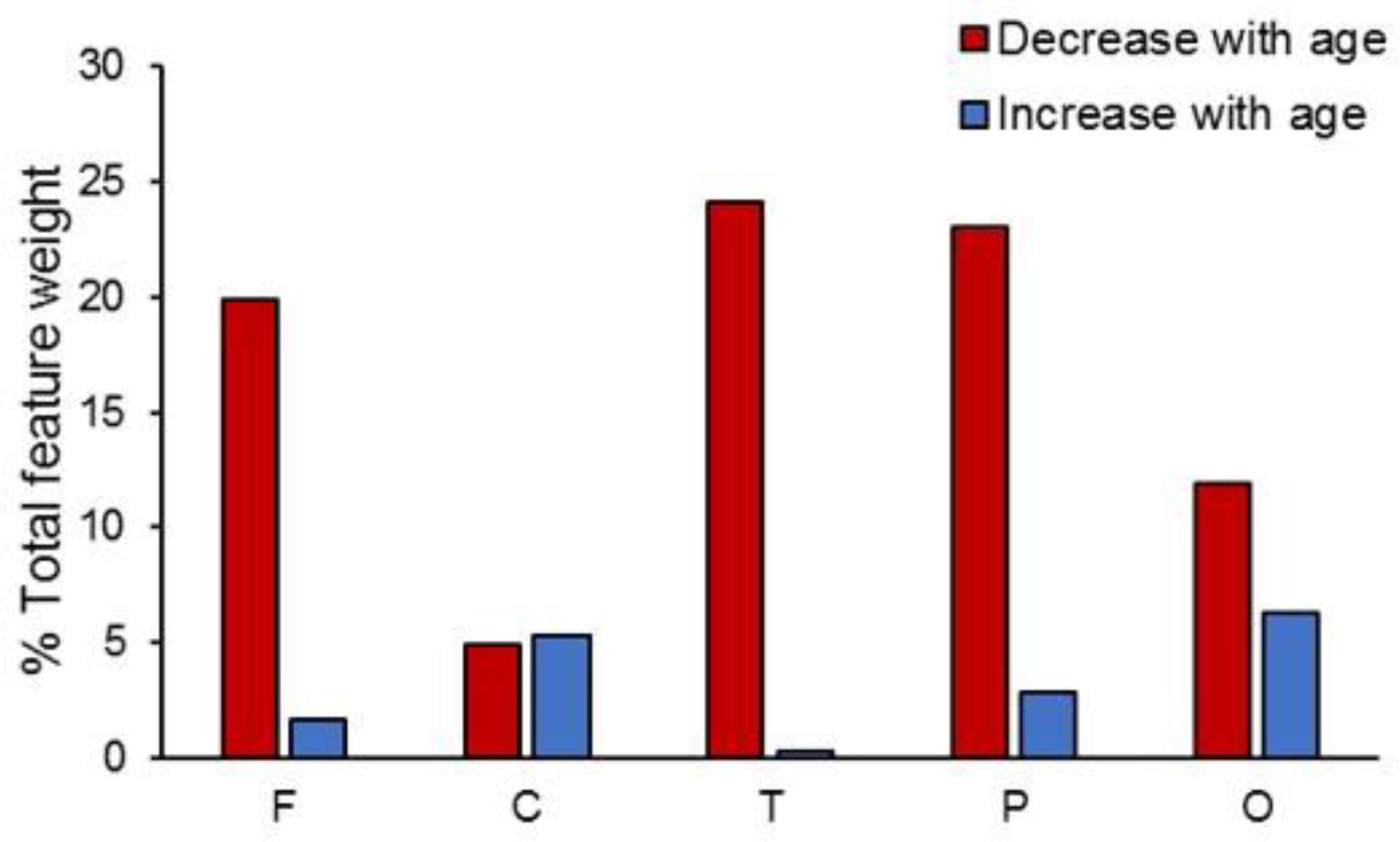
Comparison of the contribution of electrodes from each region to the SVM classification (expressed as % of total feature weight). Values are corrected for the total number of electrodes included in each region.

#### 3.4.3 Distance

We compared the connections that decreased with older age to those that increased with older age based on electrode-to-electrode Euclidian distance. We observed that consensus features showing decreased functional connectivity with older age were longer than those that showed increased connectivity with older age (*U* = 1289, *Z* = −2.6, *p* = 0.009; decrease: median = 140.9 mm, range = 29.4-185.4 mm; increase: median = 120.8 mm, range = 36.8-156.0 mm) – see Figure 5.

**Figure 5.**
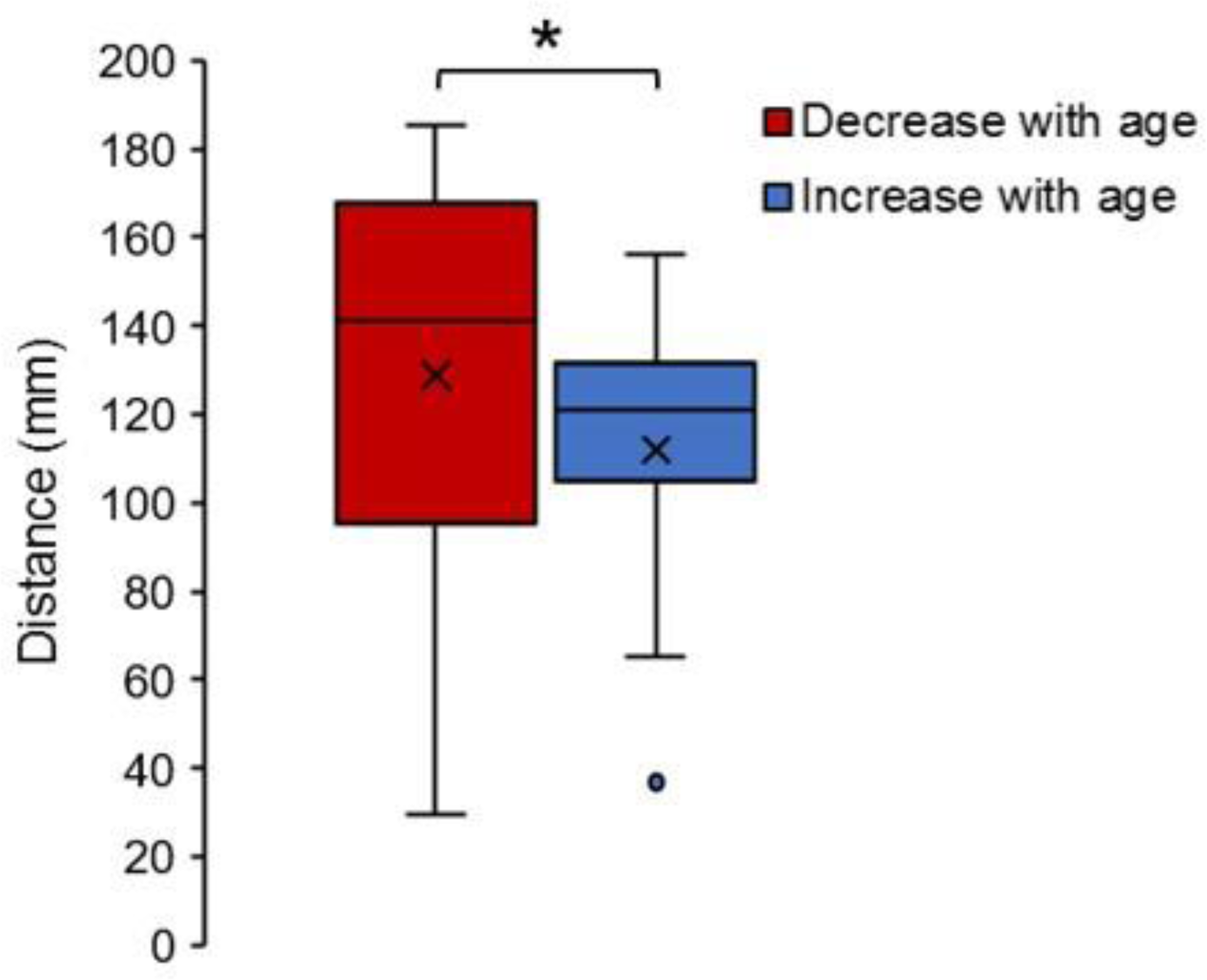
Comparison of consensus features with decreased weight with older age to those with increased weight with older age based on electrode-to-electrode distance. Box plots showing the mean (cross), median, interquartile and range of electrode-to-electrode distances for consensus features with decreased weight with older age (red) and those with increased weight with older age (blue). \**p* < 0.01.

## 4. Discussion

The greatest contribution to the classification of young from old brains came from a decrease in functional connectivity of electrodes approximating the frontal, parietal and temporal areas in older compared to young adults, particularly in alpha band. We also observed an age-related increase in functional connectivity of electrodes approximating the motor regions in the beta band. Our findings suggest a decrease in connectivity in key networks and frequency bands associated with attention and an increase in the networks and frequency band associated with motor control with advanced ageing

Ageing is characterized by alterations in functional brain networks (Ferreira & Busatto, 2013; Hedden & Gabrieli, 2004; Raz & Rodrigue, 2006; Rossini, Rossi, Babiloni, & Polich, 2007). Resting-state functional connectivity has shown to commonly decrease in healthy aging (for a review see Ferreira and Busatto (2013)). Most commonly, using fMRI decreases in frontal and parietal areas with older ageing have been observed (Hafkemeijer et al., 2012; Mevel et al., 2011). In line with this, we found that the most influential connections for classifier performance were those showing decreased connectivity with older age. This was largely due to connections involving frontal, temporal, parietal, and (to a lesser extent) occipital electrodes, with the greatest contribution coming from decreased connectivity in the alpha frequency band.

The decreased alpha band connectivity of frontal and parietal areas in older relative to younger adults may reflect changes in attentional and arousal levels with advanced ageing (Mathewson et al., 2012), with theta and alpha band oscillations suggested to reflect attentional load and arousal level (Klimesch, 2012; Klimesch, Doppelmayr, Russegger, Pachinger, & Schwaiger, 1998; Mathewson et al., 2012). Using EEG and MEG recordings, decreased alpha connectivity of frontal, parietal and temporal regions have been observed in pathological mild cognitive impairment and dementia (Babiloni et al., 2018; Babiloni et al., 2006; Babiloni et al., 2004; Bosboom, Stoffers, Wolters, Stam, & Berendse, 2009). A frontoparietal network is often engaged during tasks requiring attention (Bonnelle et al., 2011; Coull, Frackowiak, & Frith, 1998) so a decrease in frontoparietal connectivity is likely to reflect impaired attention.

Whereas consensus features for alpha, theta, and gamma bands all showed decreased functional connectivity with older age, features for beta showed increased connectivity in older relative to younger adults, accounting for 17% of the total feature weight. Neural oscillations in the beta frequency band have been strongly linked to sensorimotor network activity (Pfurtscheller et al., 1996; Roopun et al., 2006), with resting-state beta connectivity associated with motor cortex function and plasticity in healthy subjects (Hordacre et al., 2017; J. Wu, Srinivasan, Kaur, & Cramer, 2014) and after stroke (J. Wu et al., 2015). The beta band features in this study mostly involved central electrodes overlying sensorimotor networks. FMRI studies have shown age-related increases in functional connectivity of sensorimotor networks both during task performance (Heitger et al., 2013) and at rest (Langan et al., 2010; Meier et al., 2012; Mowinckel, Espeseth, & Westlye, 2012; Solesio-Jofre et al., 2014; Tomasi & Volkow, 2012), reflecting a loss of functional segregation among sensorimotor areas that is related to poorer performance on motor coordination tasks (Solesio-Jofre et al., 2014). While less is known about the role of resting-state beta rhythms in ageing, there is some evidence that advanced age is associated with increases in both resting-state beta power and beta de-synchronization during movement, possibly reflecting changes in the balance between GABAergic inhibition and glutamatergic excitation in the aged motor cortex (Rossiter, Davis, Clark, Boudrias, & Ward, 2014). Our results showing increased beta band functional connectivity and decreased theta and alpha band functional connectivity with older age are in agreement with previous research (Vysata et al., 2014). While age differences in excitatory-inhibitory balance might have contributed to age-related increases in beta band functional connectivity to classifier performance in the present study (Rossiter et al., 2014), additional research is needed to explore this further.

We observed that consensus features showing decreased functional connectivity with older age were significantly longer than those that showed increased connectivity with older age. This is in agreement with the findings of Meier et al. (2012), who reported a decrease in long-range functional connectivity and an increase in short-range functional connectivity with older age. While the physiological significance of distances separating surface electrode pairs should be interpreted with caution, these data are consistent with the notion that long-range connections, particularly between anterior and posterior brain regions, are more affected than short-range connections in normal and pathological ageing (Sala-Llonch et al., 2014; Tomasi & Volkow, 2012).

There are several limitations to this study. First, we acknowledge that EEG suffers from poor spatial resolution and signals that are recorded at the surface level can be affected by volume conduction (Bastos & Schoffelen, 2016). We have tried to mitigate this effect by using a conservative measure of functional connectivity (i.e. imaginary coherence). However, this may not thoroughly remove the effects of volume conduction and caution is needed when suggesting generators of the neural signal recorded with EEG surface electrodes. Second, the use of SVM requires several user decisions. For example, for estimating parameters we had to choose an appropriate kernel and its parameters. Because the data was not linearly separable, we chose a radial RBF kernel which projects the data to infinite dimensions and linearly separates both the groups (Meier et al., 2012). The parameters for the kernel are estimated using the nested leave one out cross-validation procedure. Finally, as mentioned above, it is important to note that using the Euclidian distance between electrodes does not reflect the distance of physical connections.

In summary, we show that an individual resting-state EEG recording can be used to classify younger adult brains from older adult brains with high accuracy, enabling greater understanding of functional connectivity as a mechanism mediating changes in human behaviour with ageing. Our study supports the literature that suggest functional connectivity of frontal, parietal and temporal regions decreases, and connectivity of motor areas increases with older age. These findings also emphasize the role of alpha frequency band connectivity in characterizing advanced ageing.

## Acknowledgements

MRG is supported by a NHMRC-ARC Dementia Research Development Fellowship (1102272). HADK is supported by a NHMRC Boosting Dementia Research Leadership Fellowship (GNT1135676). BH funded by a NHMRC fellowship (1125054). CB is funded by a NHMRC fellowship (1127155). BM would like to thank Luke Hallam for helpful discussions. We would like to thank all of the participants that made this work possible.

## Supplementary material

**Table 2.** Information on consensus features (FDR corrected for multiple comparison, *q* < 0.05). Positive features denote decreased functional connectivity with older age and negative features denote increased functional connectivity with older age.

| Frequency | Electrode 1 | Electrode 2 | Feature weights | P-value |
| --- | --- | --- | --- | --- |
| Alpha | T7 | F8 | 4.79 | 0.0008 |
| Alpha | P2 | P5 | 4.78 | 0.0008 |
| Beta | C2 | O2 | -4.74 | 0.0008 |
| Beta | PO8 | C2 | -4.71 | 0.0008 |
| Alpha | PO3 | P2 | 4.61 | 0.0008 |
| Alpha | TP7 | AF4 | 4.55 | 0.0008 |
| Alpha | P1 | AF4 | 4.54 | 0.0008 |
| Alpha | PO5 | POz | 4.48 | 0.0008 |
| Beta | PO6 | C2 | -4.44 | 0.0008 |
| Alpha | PO5 | P2 | 4.42 | 0.0008 |
| Beta | Oz | C3 | -4.40 | 0.0008 |
| Alpha | TP7 | F8 | 4.35 | 0.0008 |
| Alpha | PO3 | Pz | 4.34 | 0.0008 |
| Alpha | POz | F4 | 4.31 | 0.0008 |
| Alpha | P2 | T8 | 4.31 | 0.0008 |
| Alpha | PO3 | POz | 4.31 | 0.0008 |
| Alpha | TP7 | F4 | 4.30 | 0.0008 |
| Alpha | TP7 | P7 | 4.30 | 0.0008 |
| Alpha | P5 | POz | 4.28 | 0.0008 |
| Alpha | TP7 | P1 | 4.26 | 0.0008 |
| Alpha | FT7 | Pz | 4.25 | 0.0008 |
| Alpha | FT7 | P2 | 4.24 | 0.0008 |
| Alpha | TP7 | AF3 | 4.23 | 0.0008 |
| Alpha | FT7 | CP2 | 4.19 | 0.0009 |
| Alpha | P7 | T7 | 4.17 | 0.0010 |
| Alpha | P5 | AF4 | 4.16 | 0.0010 |
| Alpha | CP4 | AF3 | 4.14 | 0.0010 |
| Alpha | POz | FC6 | 4.13 | 0.0010 |
| Beta | C2 | Oz | -4.12 | 0.0010 |
| Alpha | P4 | CP6 | 4.11 | 0.0010 |
| Alpha | P5 | Pz | 4.09 | 0.0010 |
| Gamma | CP3 | CP1 | 4.07 | 0.0011 |
| Alpha | P1 | F8 | 4.06 | 0.0011 |
| Alpha | TP7 | Pz | 4.06 | 0.0011 |
| Gamma | CP4 | F5 | 4.02 | 0.0012 |
| Alpha | TP7 | P5 | 4.02 | 0.0012 |
| alpha | TP7 | Fz | 4.00 | 0.0012 |
| alpha | PO7 | T7 | 4.00 | 0.0012 |
| alpha | PO7 | POz | 3.99 | 0.0012 |
| alpha | CP2 | T7 | 3.97 | 0.0012 |
| Beta | PO4 | C2 | -3.97 | 0.0012 |
| Alpha | P2 | P3 | 3.95 | 0.0012 |
| Alpha | TP7 | F3 | 3.95 | 0.0012 |
| Alpha | P2 | CP5 | 3.95 | 0.0012 |
| Alpha | P5 | P4 | 3.90 | 0.0014 |
| Alpha | TP7 | FC1 | 3.89 | 0.0014 |
| Alpha | FC2 | POz | 3.88 | 0.0014 |
| Alpha | TP7 | FC2 | 3.88 | 0.0014 |
| Alpha | TP7 | FC6 | 3.88 | 0.0014 |
| Alpha | FC4 | POz | 3.87 | 0.0014 |
| Alpha | Pz | CP5 | 3.86 | 0.0014 |
| Alpha | CP4 | F7 | 3.86 | 0.0014 |
| Beta | CPz | O2 | -3.85 | 0.0014 |
| Alpha | P2 | F7 | 3.85 | 0.0014 |
| Alpha | PO7 | P2 | 3.84 | 0.0014 |
| Alpha | P1 | T7 | 3.83 | 0.0014 |
| Alpha | AF4 | Pz | 3.83 | 0.0014 |
| Alpha | CP4 | F5 | 3.83 | 0.0014 |
| Alpha | TP7 | PO5 | 3.82 | 0.0014 |
| Gamma | F5 | CP6 | 3.82 | 0.0014 |
| Alpha | Pz | T7 | 3.81 | 0.0014 |
| Alpha | P2 | F3 | 3.81 | 0.0014 |
| Alpha | TP7 | FC6 | 3.79 | 0.0014 |
| Beta | C2 | Cz | -3.76 | 0.0015 |
| Alpha | TP7 | FC2 | 3.76 | 0.0015 |
| Alpha | FC1 | 17CP6 | 3.76 | 0.0015 |
| Alpha | PO5 | Pz | 3.75 | 0.0016 |
| Alpha | Pz | F7 | 3.75 | 0.0016 |
| Theta | T7 | Fz | 3.72 | 0.0017 |
| Theta | C5 | CP5 | 3.72 | 0.0017 |
| Beta | C5 | F5 | -3.71 | 0.0017 |
| alpha | P3 | T7 | 3.71 | 0.0017 |
| alpha | FC6 | T7 | 3.71 | 0.0017 |
| Theta | TP7 | Fz | 3.70 | 0.0017 |
| alpha | FC2 | T7 | 3.70 | 0.0017 |
| alpha | TP7 | P3 | 3.69 | 0.0017 |
| alpha | P1 | CP5 | 3.68 | 0.0017 |
| Theta | T8 | Fz | 3.68 | 0.0017 |
| Beta | O1 | C3 | -3.67 | 0.0017 |
| Alpha | Pz | P3 | 3.66 | 0.0018 |
| Alpha | P3 | CP5 | 3.66 | 0.0018 |
| Alpha | C4 | T7 | 3.63 | 0.0019 |
| Alpha | P7 | F7 | 3.63 | 0.0019 |
| Alpha | TP7 | C4 | 3.63 | 0.0019 |
| Alpha | T7 | FC2 | 3.62 | 0.0019 |
| Alpha | FC1 | P4 | 3.62 | 0.0019 |
| Alpha | P5 | F7 | 3.62 | 0.0019 |
| Alpha | CP2 | F7 | 3.61 | 0.0019 |
| Alpha | PO7 | TP7 | 3.61 | 0.0019 |
| Alpha | FT7 | F4 | 3.61 | 0.0019 |
| Gamma | T7 | F8 | 3.61 | 0.0019 |
| Beta | C1 | O1 | -3.60 | 0.0019 |
| Alpha | AF3 | P7 | 3.60 | 0.0019 |
| Alpha | CP5 | F8 | 3.59 | 0.0019 |
| Gamma | FC1 | F3 | 3.59 | 0.0019 |
| Alpha | AF3 | CP6 | 3.58 | 0.0019 |
| Alpha | CPz | T7 | 3.58 | 0.0019 |
| Alpha | P1 | Pz | 3.57 | 0.0020 |
| Alpha | Pz | FC6 | 3.56 | 0.0020 |
| Theta | PO8 | FC1 | 3.55 | 0.0021 |
| Alpha | AF3 | P4 | 3.55 | 0.0021 |
| Theta | C5 | FC1 | 3.54 | 0.0021 |
| Alpha | AF4 | P3 | 3.53 | 0.0022 |
| Alpha | PO7 | FC1 | 3.50 | 0.0024 |
| Alpha | TP7 | FC4 | 3.49 | 0.0024 |
| Alpha | PO5 | T7 | 3.49 | 0.0024 |
| Alpha | AF4 | P4 | 3.48 | 0.0025 |
| Alpha | C2 | T7 | 3.47 | 0.0025 |
| Theta | TP7 | FC1 | 3.46 | 0.0025 |
| Beta | O2 | C4 | -3.46 | 0.0025 |
| Alpha | FT8 | CPz | 3.46 | 0.0025 |
| Theta | FC2 | T8 | 3.45 | 0.0025 |
| Alpha | O2 | POz | 3.45 | 0.0025 |
| Theta | PO4 | T7 | 3.45 | 0.0025 |
| Beta | C1 | Oz | -3.45 | 0.0025 |
| Beta | Oz | CP1 | -3.43 | 0.0027 |
| Alpha | FC2 | FC6 | 3.42 | 0.0027 |
| Beta | PO7 | Pz | -3.42 | 0.0027 |
| Alpha | PO5 | P4 | 3.42 | 0.0027 |
| Alpha | PO8 | POz | 3.42 | 0.0027 |
| Alpha | P1 | C5 | 3.41 | 0.0027 |
| Alpha | AF4 | CP6 | 3.41 | 0.0027 |
| Theta | FC1 | CP5 | 3.41 | 0.0027 |
| Beta | CPz | Oz | -3.40 | 0.0027 |
| Alpha | TP7 | C6 | 3.38 | 0.0028 |
| Alpha | Pz | F8 | 3.37 | 0.0029 |
| Alpha | PO5 | F7 | 3.37 | 0.0030 |
| Alpha | P2 | AF3 | 3.35 | 0.0031 |
| Beta | F5 | Cz | -3.35 | 0.0031 |
| Alpha | PO3 | P4 | 3.34 | 0.0031 |
| Alpha | P2 | CP6 | 3.33 | 0.0032 |
| Theta | PO7 | TP7 | 3.33 | 0.0032 |
| Alpha | CP4 | C6 | 3.32 | 0.0032 |
| Alpha | P4 | F3 | 3.32 | 0.0033 |
| Alpha | F5 | Pz | 3.31 | 0.0033 |
| Beta | O2 | CP1 | -3.30 | 0.0034 |
| Alpha | FC2 | Pz | 3.30 | 0.0034 |
| Beta | CPz | CP1 | -3.29 | 0.0035 |
| Theta | FT8 | Fz | 3.28 | 0.0035 |
| Beta | C2 | F7 | -3.28 | 0.0035 |
| Beta | PO8 | CPz | -3.27 | 0.0036 |
| Beta | O2 | CP2 | -3.26 | 0.0037 |
| Beta | C1 | AF4 | -3.25 | 0.0037 |
| Alpha | Pz | Fz | 3.25 | 0.0037 |
| Alpha | PO4 | P5 | 3.24 | 0.0038 |
| Beta | FT7 | FC2 | -3.24 | 0.0038 |
| Alpha | C6 | P4 | 3.24 | 0.0038 |
| Alpha | FT8 | P2 | 3.23 | 0.0039 |
| Alpha | FT7 | C4 | 3.22 | 0.0040 |
| Alpha | C5 | FC6 | 3.21 | 0.0041 |
| Alpha | TP7 | P2 | 3.21 | 0.0041 |
| Beta | C2 | C1 | -3.20 | 0.0041 |
| Beta | FC6 | Cz | -3.20 | 0.0042 |
| Beta | C5 | F3 | -3.19 | 0.0043 |
| Theta | F8 | Fz | 3.18 | 0.0043 |
| Beta | O1 | Cz | -3.17 | 0.0044 |
| Alpha | FT7 | P4 | 3.17 | 0.0044 |
| Theta | TP7 | O1 | 3.15 | 0.0046 |
| Theta | FC1 | T7 | 3.13 | 0.0050 |
| Theta | FC2 | T7 | 3.11 | 0.0052 |
| Theta | FT7 | POz | 3.05 | 0.0061 |
| Alpha | PO4 | Fz | 3.02 | 0.0068 |

